# Clonal hematopoiesis detection in cancer patients using cell free DNA sequencing

**DOI:** 10.1101/2021.10.27.466159

**Authors:** Lauren Fairchild, Jeanne Whalen, Katie D’Aco, Jincheng Wu, Carroll B. Gustafson, Nadia Solovieff, Fei Su, Rebecca Leary, Catarina D. Campbell, O. Alejandro Balbin

**Author notes:** These authors jointly supervised this work (Catarina D. Campbell, O. Alejandro Balbin).

## Abstract

In the context of cancer, clonal hematopoiesis of indeterminate potential (CHIP) is associated with the development of therapy-related myeloid neoplasms and shorter overall survival. Cell-free DNA (cfDNA) sequencing is becoming widely adopted for genomic screening of cancer patients but has not been used extensively to determine CHIP status due to a requirement for matched blood and tumor sequencing. Here we present an accurate machine learning approach to determine clonal hematopoiesis (CH) status from cfDNA sequencing alone and apply our model to 4,096 oncology clinical cfDNA samples. Using this method, we determine that 26% of patients in this cohort have evidence of CH and CH is most common in lung cancer patients. Matched RNAseq data shows signals of increased inflammation, especially neutrophil activation, within the tumor microenvironment of CH-positive patients. Additionally, CH patients showed evidence of neutrophil activation systemically, pointing to a potential mechanism of action for the worse outcomes associated with CH status. Neutrophil activation may be one of many mechanisms however, as estrogen positive breast cancer patients harboring TET2 frameshift mutations had worse outcomes but similar neutrophil levels to CH-negative patients.

**One Sentence Summary:** We train an accurate machine learning model to detect clonal hematopoiesis in cancer and characterize associated changes in the tumor microenvironment.

## INTRODUCTION

Clonal hematopoiesis of indeterminate potential (CHIP) is a common age-associated phenotype found in about 10% of healthy individuals over 70 years of age(*1*) and is characterized by clonal expansion of hematopoietic stem cells (HSCs) or other blood progenitor cells in the absence of an overt malignancy. Despite this, the presence of CHIP has been associated with numerous adverse health outcomes, including increased risk for cardiac events and progression to overt blood malignancies such as acute myeloid leukemia (AML). CHIP and therapy-related myeloproliferative neoplasms (tMNs) are seen at higher rates in oncology patients, and CHIP is associated with poorer overall survival in the oncology setting.(*2*)

In seminal studies analyzing exome data from individuals without cancer, CHIP was defined by the presence of mutations in recognized hematological neoplasm driver genes (most commonly DNMT3A, TET2, ASXL1, JAK2, PPM1D, and SF3B1) with variant allele frequency (VAF) of >2% in the absence of severe cytopenia or other overt malignancies.(*3*) However, with increasing sequencing depth, somatic variants in these same genes have been detected at much lower VAFs in ‘healthy’ individuals.(*4*) However, the clinical impact of these low allele fraction variants is not well understood.

In an oncology setting of patients with advanced-stage solid cancers, *Bolton et al*(*5*) showed that therapy could influence the clonal dynamics of existing CHIP mutations, in addition to inducing new CHIP mutations. Specifically, cytotoxic and radiation therapy increased the size of clones with mutations in DNA damage repair pathway, leading to the hypothesis that therapy mediated neoplasms could occur as a result of selective pressures on pre-existing CHIP variants.

Cell-free DNA (cfDNA) sequencing is gaining wide usage in the clinical trial setting as a non-invasive approach for determining the genomic landscape of cancer, monitoring minimal residual disease, and potential early cancer detection. As cfDNA is a mixture of circulating normal and tumor DNA, the biological source of a mutation detected in plasma can only be determined unambiguously by comparing matched sequencing of equal depth from white blood cells (WBC) and plasma. This approach is cost-prohibitive and commercial cfDNA assays do not routinely incorporate WBC sequencing. Therefore, developing computational approaches to annotate clonal hematopoiesis (CH) variants in cfDNA sequencing data would alleviate the requirement of matched WBC sequencing data, augment the value of plasma sequencing and enable investigation into the impact of CH on cancer trajectory and progression. Furthermore, being able to make this determination from plasma sequencing alone would enable retroactive studies of CH where only cfDNA was sequenced.

Here, we present an accurate machine learning approach able to discriminate between blood-derived and tumor-derived mutations in plasma cfDNA from oncology patients without using WBC matched sequencing. We then apply this method to over 4,000 cfDNA baseline plasma samples from patients with advanced cancer to characterize CH in an oncology setting and correlate patient CH status with other peripheral and tumor microenvironment (TME) biomarkers. Our analyses suggest that CH-positive patients have increased neutrophil and inflammatory activity in the TME. This is also reflected in increases in peripheral blood counts associated with systemic inflammation, such as absolute levels of monocyte and neutrophil counts. Furthermore, CH-positive status was associated with elevated neutrophil to lymphocyte ratio (NLR) and this signal was found predominantly in estrogen receptor positive breast cancer and melanoma patients.

## RESULTS

### Subhead 1: A machine learning approach to discriminate blood-derived from tumor-derived single nucleotide variants in cfDNA

In order to establish a ground truth dataset of variants detected in cfDNA and their known biological sources, we identified a published dataset containing matched WBC, plasma, and tumor sequencing from 124 metastatic cancer patients and 47 healthy controls.(*6*) (Table S1) Using the 1,386 single nucleotide variants (SNVs) in this dataset unambiguously derived from either tumor or blood, we trained random forest and logistic regression models to classify the source of these variants using plasma data alone (Fig. 1A). Features provided to the models included variant allele frequency, median allele frequency across all detected variants in the patient, and whether the variant occurred in a canonical CHIP gene. We expanded the variant annotations by including the frequency of the variant in public datasets including COSMIC(*7*), GnomAD(*8*), and ExAC(*9*) in order to inform the model about the incidence of each variant in other cancer and healthy populations, since a variant seen often in cancer populations would be more likely to be tumor derived. Finally, we also calculated the 60 single base substitution (SBS) signatures(*10*) for each variant, as certain mutational signatures reflect environments associated with CH (aging) or with cancer (oncologic therapy). (Methods, Table S2) Due to the limited number of insertions and deletions in this dataset that could be used for training and testing machine learning algorithms, likely CH insertions and deletions were classified using a rule-based approach as described by *Jaiswal et al 2014* and *Svensson et al 2018 (11, 12)*.

**Fig. 1.**
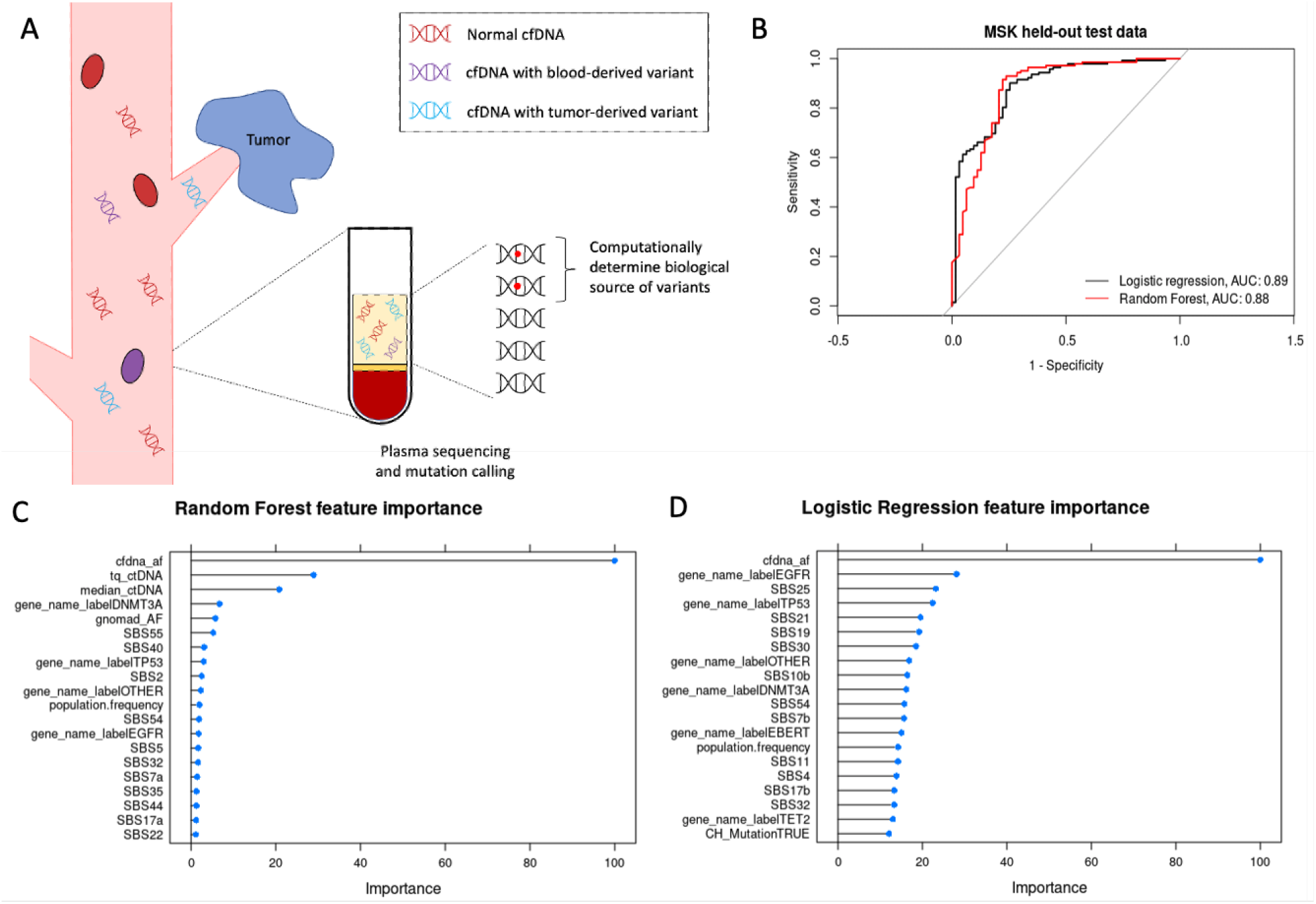
Machine learning approach and performance discriminating blood from tumor derived variants. **(A)** Variants found in both blood and tumor cells contribute to the variants found in cfDNA (plasma). It is difficult to determine the source of variants in cfDNA without matched blood and tumor sequencing. **(B)** ROC indicating performance of two machine learning approaches on a held-out test set. **(C, D)** Relative importance of features used in the random forest and logistic regression models, respectively.

Both random forest (RF) and logistic regression (LR) models were trained using a 10-fold cross validation paradigm with 15% of the data held out for testing. Both models performed well on the held-out test set, with 0.88 and 0.89 area under the receiver operator characteristic (auROC) and 0.91 and 0.90 F1 scores for the random forest and logistic regression, respectively (Figure 1B). Several methods were tested to rebalance the classes(*13, 14*), but they did not improve performance of the models, likely because the classes were not strongly imbalanced (2:1). Although the two approaches performed similarly when evaluated by auROC and F1 scores, in an additional test in which SNVs from healthy controls were held out from training and tested separately (N=243 blood-derived variants), the logistic regression significantly outperformed the random forest with 13 misclassified control variants versus 29 (94.7% correct vs 88.1%, respectively, Table S3). Additionally, 8 of the 29 errors made by the random forest occurred in DNMT3A or TET2, whereas all the errors made by the logistic regression occurred in different genes. The logistic regression approach also placed importance on a broader set of features, including relevant SBS mutational signatures, than the random forest approach.

Interestingly, several of the most informative mutational signatures for the logistic regression were associated with various therapies including SBS25 (chemotherapy treatment), SBS11 (temozolomide treatment), and SBS32 (azathioprine treatment). In addition, several signatures associated with issues in DNA repair (SBS21, SBS30, SBS10b) and a signature associated with smoking (SBS4) were important in the model. In contrast, only three features were used predominantly by the random forest, all of which were measures of allele fraction. (Figure 1C, D)

### Subhead 2: Application of CH classification method to oncology clinical samples

To further test our models and to investigate the role of clonal hematopoiesis in cancer, we curated a set of baseline cfDNA sequencing data from 4,296 advanced metastatic cancer patients enrolled in Novartis oncology clinical trials. Patients were predominantly diagnosed with estrogen receptor positive breast cancer (BRCA, 56.4%), cutaneous melanoma (13.6%), non-small cell lung cancer (NSCLC, 11.4%), colorectal cancer (CRC, 5.7%), or triple negative breast cancer (TNBC, 4.7%) (Table 1). Data was generated using two versions of a targeted DNA sequencing panel of about 564 genes (Methods). Although the lower limit of detection varied slightly between the two versions of this panel, there was no difference in the proportion of variants predicted to be blood-derived in canonical CHIP genes and no difference in the detection rate of CH at the patient level (Methods, Table S4).

**Table 1.** Summary of clinical data. Shown are only indications with at least 100 cfDNA samples. Other indications include anal cancer, anaplastic thyroid cancer, bladder cancer, cervical cancer, cholangiocarcinoma, chordoma, endometrial cancer, esophageal cancer, gallbladder cancer, gastric cancer, head and neck cancer, hepatocellular carcinoma, liposarcoma, malignant neoplasm of the thymus, merkel cell carcinoma, mesothelioma, nasopharyngeal cancer, neuroendocrine tumor, non-cutaneous/uveal melanoma, non-Hodgkin’s lymphoma, ovarian cancer, pancreatic cancer, prostate cancer, renal cell carcinoma, sarcoma, squamous cell carcinoma, testicular cancer, thyroid cancer.

| Tumor type | Baseline<br>cfDNA | Matched<br>baseline RNA | Median age of<br>cohort (years) |
| --- | --- | --- | --- |
| ER<sup>+</sup> Breast cancer | 2423 | 103 | 60 |
| Cutaneous melanoma | 585 | 43 | 56 |
| Non-small cell lung cancer | 491 | 130 | 63 |
| Colorectal cancer | 248 | 190 | 59 |
| Triple negative breast cancer | 204 | 157 | 52 |
| Other or Unknown | 343 | 188 | - |
| Total | 4294 | 795 | - |

Both the LR and RF models were used to classify the 34,855 variants detected in this internal dataset. The logistic regression labeled more variants as blood-derived than the random forest (26% vs 17%). We split the predicted blood-derived variants into those within likely CHIP driver genes(*11*) (DNMT3A, TET2, ASXL1, JAK2, SF3B1, and PPM1D, 15.7%) and those within other frequently observed myeloid or hematological malignancy driver genes(*5*) (19.4%, Fig. 2A, Methods, Table S5). The genes with the most predicted blood-derived variants from both approaches were DNMT3A and TET2, as expected. Other less common CHIP genes such as SF3B1, CBL, and KMT2C were also enriched for blood-derived variants. The LR approach classified many more TET2, SF3B1, and CBL variants as blood derived, relative to random forest, and called fewer blood-derived variants in known oncogenes KRAS and EGFR (Figs. 2B, S1). For these reasons, as well as the performance on the published data, we concluded that the logistic regression model was better able to classify SNVs as blood-derived or tumor-derived and we used this approach for the remainder of our analysis.

**Fig. 2.**
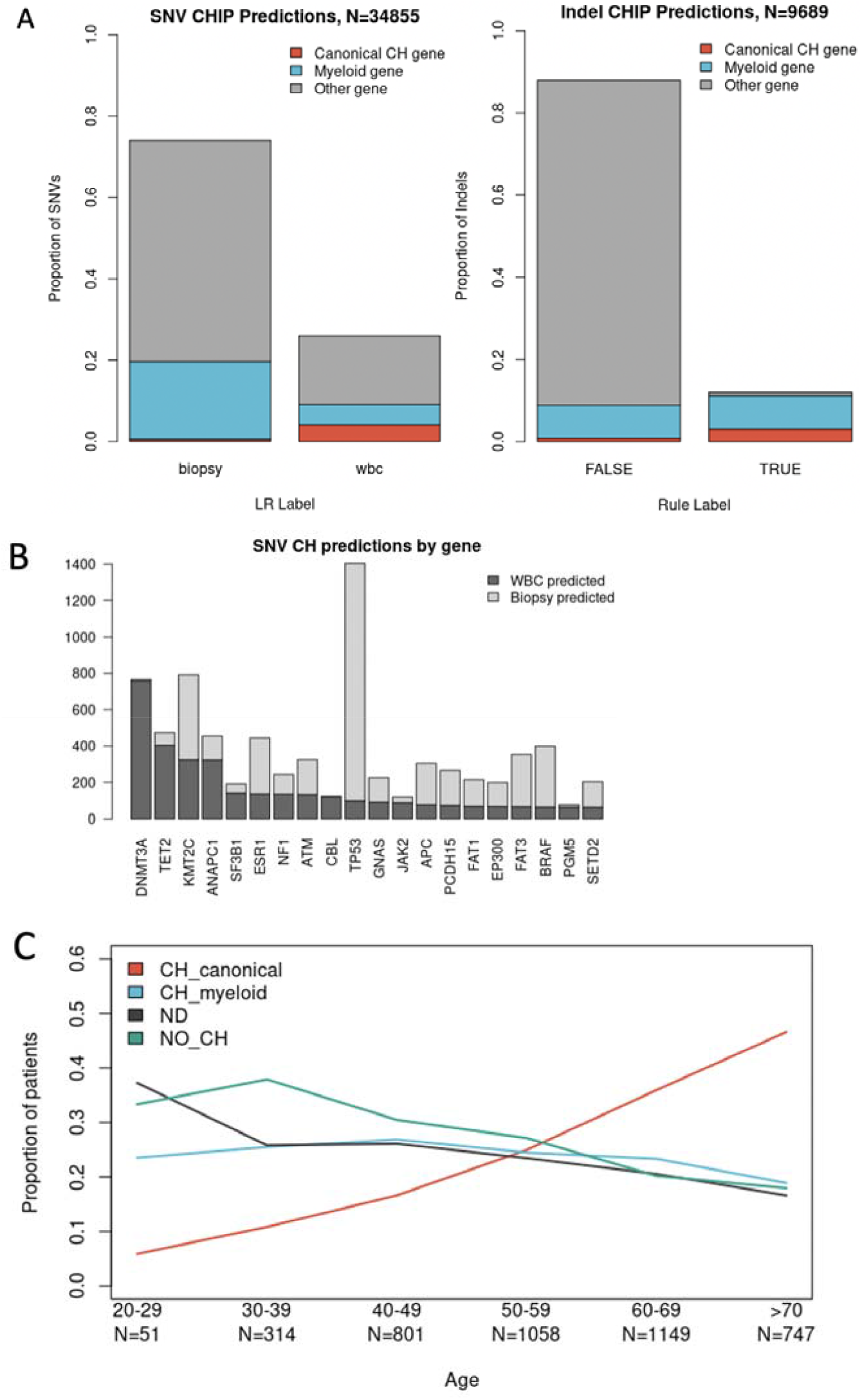
Characterization of CH in oncology patients. (**A**) Proportion of SNVs classified by the logistic regression approach (left) and indels classified with pre-defined rules (right) as either biopsy-derived or WBC-derived. Colors indicate whether the variant appeared in a canonical CHIP gene (orange), myeloid-driver gene (blue), or other gene (grey). (**B**) Proportion of SNVs within each gene classified by logistic regression as WBC-derived (dark grey) vs biopsy-derived (light grey). (**C**) Proportion of patients in each CH category plotted by age.

In order to determine CH status at a patient level, we combined the SNV CH predictions from the logistic regression approach (regardless of VAF) with CH labels for indels derived using a rule-based approach (Methods)(*12*). A patient was labeled CH-positive if one or more blood-derived variants was present in the pre-defined set of canonical CHIP genes. If a patient did not have blood-derived variants in canonical CHIP genes but a blood-derived variant was found in a putative myeloid driver gene, the patient was labeled CH-myeloid. If a patient was neither CH-positive nor CH-myeloid, the patient was labeled CH-negative (Methods). Using this framework, we found that 28.9% of patients in our dataset were CH-positive (12.9% with >2% VAF). Similar to previous reports(*11*), most patients with CH mutations had only one variant (75.0%, Fig. S2). The proportion of CH-positive patients increased substantially with age from 10.1% in patients under 40 to 46.5% in patients over 70 years of age (N=365 and 747, respectively. P-value<2.2e-16, Fisher’s exact test). The proportion of patients with blood-derived variants in the putative myeloid driver gene set remained consistent regardless of age. (Fig. 2C) In comparison with healthy individuals, the proportion of CH-positive cancer patients found here is higher, agreeing with previous reports of an increased incidence of CH in oncology cohorts (Fig. S3)(*1, 5, 11*).

We next sought to determine if there was any difference in the incidence of CH in patients with different tumor types. Using a logistic regression model to account for differences in age between cohorts, we found that NSCLC patients were significantly more likely to have CH mutations than patients with other types of tumors (TNBC, BRCA, melanoma, CRC) with 41.8% of patients classified as CH-positive versus 26.8% in other indications. This trend remained after correcting for age (Odds-ratio: 1.77 (1.40-2.25) relative to BRCA, p-value: 1.59e-6). All other tumor types were equally likely to have CH after correcting for differences in patient age. (Fig. 3A, B) We considered the possibility that NSCLC patients had higher incidence of CH due to differences in prior therapy but including prior chemotherapy status as a covariate in the model did not change the odds of having CH for any indication (Fig S4A). Prior work by *Bolton et al(5)* showed an increase in CH in current and former smokers relative to non-smokers, and this may contribute to the increased incidence we observe in this NSCLC cohort. Unfortunately, smoking status was not curated for enough patients to confirm this hypothesis in this cohort. When repeating this analysis comparing the odds of being CH-positive or CH-myeloid versus having no evidence of blood-derived variants in either of these gene sets, NSCLC, CRC, and TNBC were all more likely to have these mutations than BRCA (Fig. S4B).

**Fig. 3.**
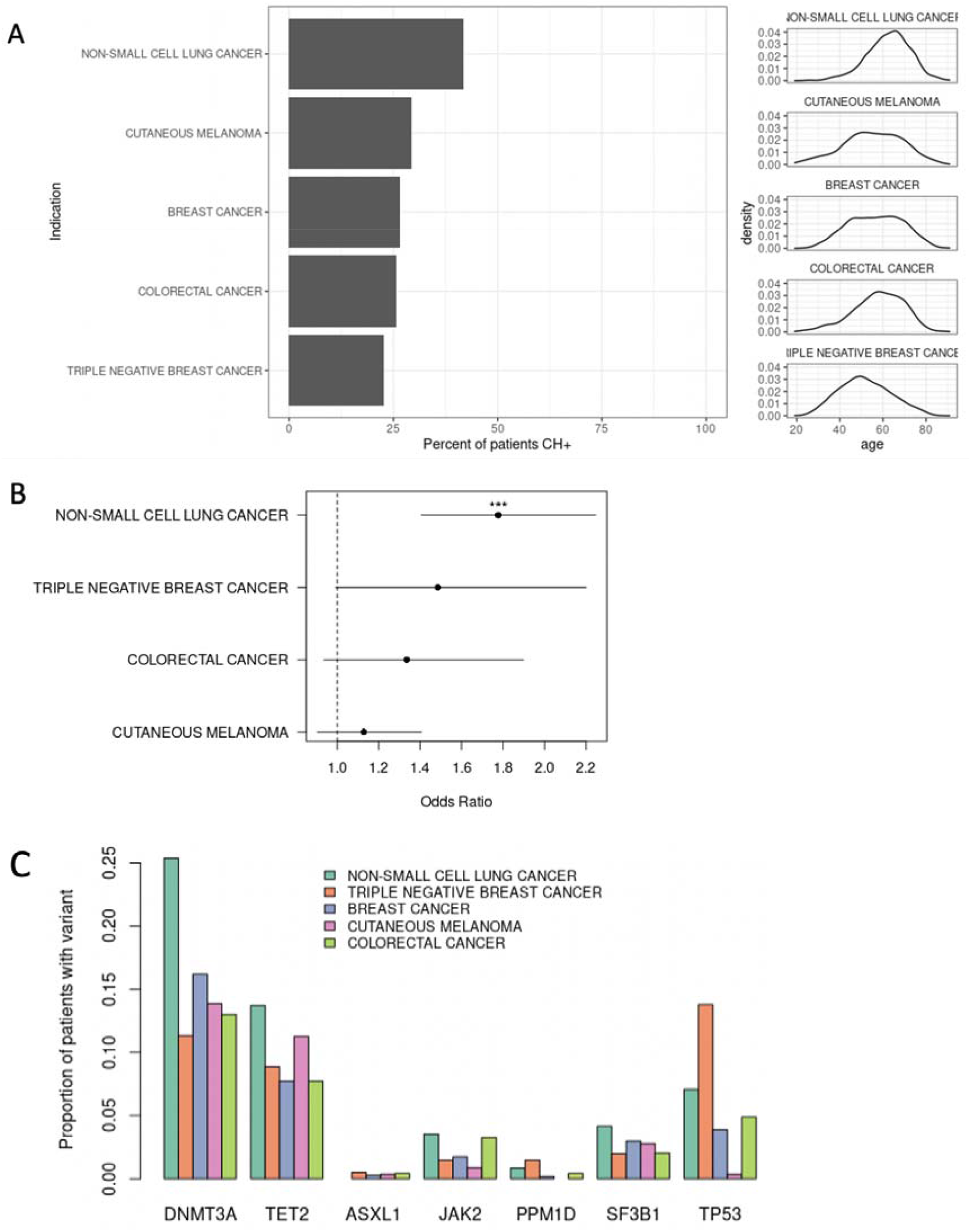
Characterization of CH in patients with advanced cancer. (**A**) Percent of patients predicted to be CH-positive split by tumor type (left) with corresponding distribution of patient ages(right) (**B**) Odds-ratio of CH by tumor type relative to breast cancer accounting for age of patients. (NSCLC p-value =2.96e-7) (**C**) Proportion of patients with mutations in common CH genes split by tumor type.

Looking more deeply, NSCLC patients were more likely than patients with other cancer types to have blood-derived mutations in DNMT3A and JAK2, whereas patients with melanoma or NSCLC were more likely to have variants in TET2. Predicted blood-derived TP53 variants were most frequent in TNBC patients (Figs. 3C, S5). ASXL1 was found at a lower rate than previously reported, but this is because the ASXL1 gene is only included in one of the two gene panels used to generate our internal dataset.

### Subhead 3: CH status is correlated with increased expression of inflammation gene signatures in the tumor microenvironment

Since CHIP status, particularly TET2 or DNMT3A mutation, has been correlated with increased systemic inflammation markers (circulating IL-6(*5, 15*) and IL-8, and increases in IL-1β, IL-6, and chemokine expression(*16, 17*)), we investigated the association of CHIP status with tumor microenvironment gene expression. To explore the gene expression differences in the TME between CH-positive and negative patients, we explored matched baseline tumor RNAseq for 811 of the 4,295 patients with cfDNA sequencing. Within this set, 255 patients (31.4%) were predicted to be CH-positive, and an additional 265 patients (32.7%) were CH-myeloid. When comparing CH-positive against CH-negative patients, we found 153 genes were up-regulated and 87 genes were down-regulated after correcting for tumor type, biopsy site, and a liver confounder score (FDR-corrected p-value < 0.05, Methods). When we performed gene ontology (GO) term enrichment analysis, we found the up-regulated genes to be significantly enriched in terms relating to neutrophil degranulation and extravasation and inflammatory response (p<0.05, Fig. 4A). This effect was seen consistently in NSCLC, BRCA and CRC, but the effect was weaker in TNBC (Fig S6). We believe this was due to the smaller number of CH-positive TNBC patients, likely due to the lower median age of the cohort (52 years) in comparison with NSCLC (63 years), ER+ BRCA (60 years), and CRC (59 years). Expression of IL-6 and mean expression of a neutrophil linage gene signature(*18*) were also up-regulated in the TME of CH-positive patients relative to CH-negative (log_2_ fold change = 0.03, 0.15, respectively). This signal was stronger in patients with >2% VAF in their CH mutations (log_2_ fold change = 0.08, 0.17, respectively), adding evidence that the effect is related to CH status (Fig. 4B).

**Fig. 4.**
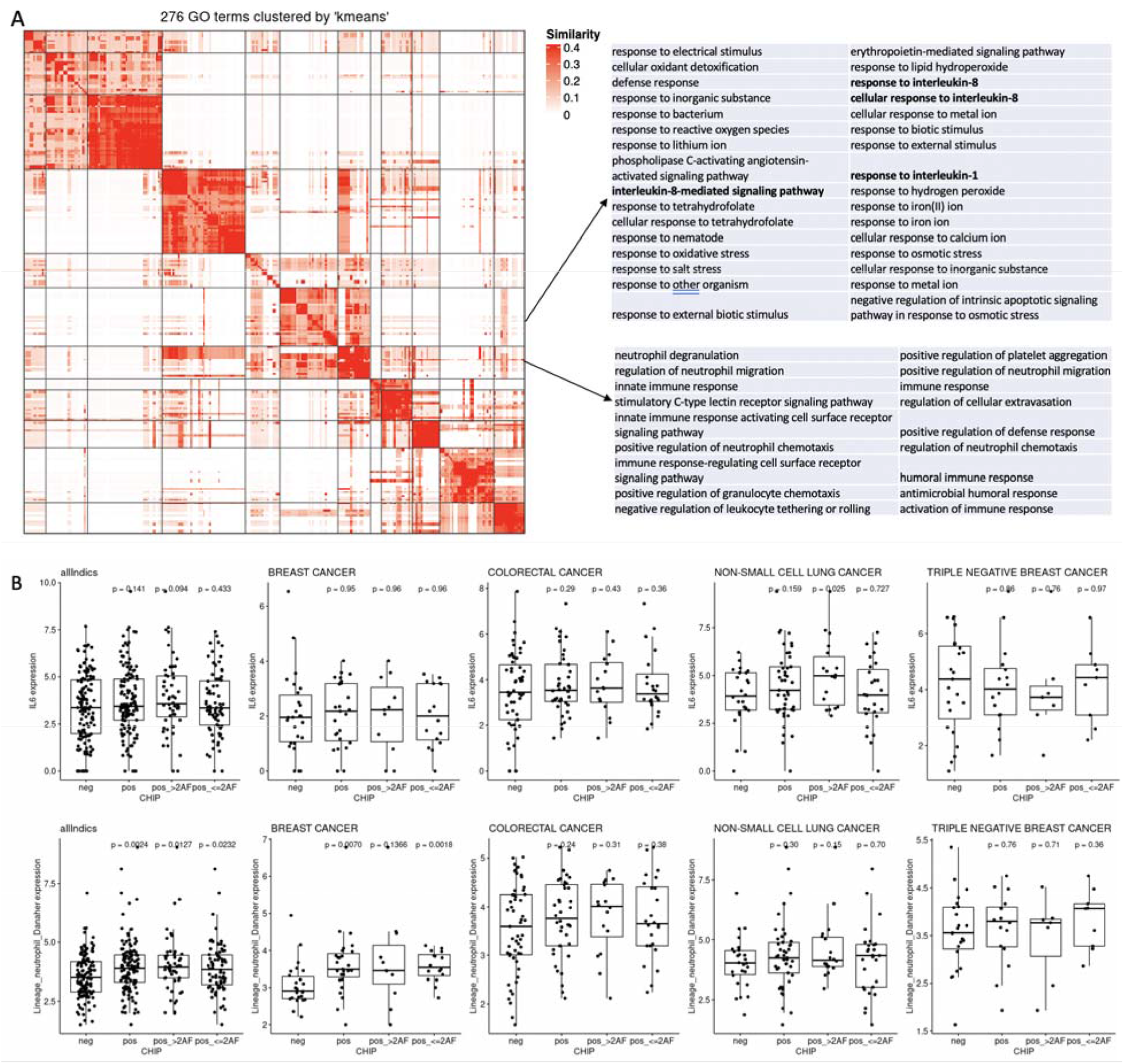
Gene expression changes in the TME related to CH status. (**A**) Significantly enriched GO terms in CH-positive patients relative to CH-negative patients clustered by GO-term gene membership similarity. Differential expression was calculated across all patients correcting by tumor type. (**B**) Expression of IL-6 and mean expression of a neutrophil lineage gene signature by tumor type (p-values calculated by T-test comparing to CHIP-negative patients, AF=allele fraction)

Finally, we wanted to determine if the observed increase in neutrophil and inflammatory activity seen in the TME of CH-positive patients, measured by RNAseq, was reflected in changes in peripheral blood counts associated with systemic inflammation. Looking at blood count data for patients classified as CH-positive, CH-myeloid, and CH-negative, we found differences in several blood analytes. Hematocrit and hemoglobin levels were lowest in CH-positive (125 g/L and 0.37 (L/L), respectively), followed by CH-myeloid (125 g/L, 0.375 L/L) and CH-negative patients (127 g/L, 0.38 L/L). Furthermore, increased absolute monocyte and neutrophil counts were noted with respect to CH status, with the highest values found in the CH-positive patient group (CH-negative: 0.4, 4. CH-myeloid: 0.43, 4.15. CH-positive: 0.45, 4.32, respectively. All units are 10e9/L. Fig. S7).

Neutrophil-to-lymphocyte ratio (NLR) is associated with poor prognosis in many cancer types(*19*) and was elevated in both CH-positive and CH-myeloid patients (Fig. S7). When using a threshold of 3 to divide patients into high and low NLR groups(*20*), we similarly found that CH-positive status was associated with elevated NLR and that this signal was found predominantly in BRCA (OR: 1.49, p-value: 0.00027, Table S6). Although age has been shown to be associated with NLR(*19, 21*), this association between CH status and NLR remained after correcting for age using a logistic regression model. We further split CH-positive patients into those with DNMT3A and TET2 mutations to determine if mutations in a specific CHIP gene were associated with increased NLR. We found that NLR was the highest in DNMT3A-mutated patients (median of 3.12 vs 2.84, p-value = 0.043 relative to wildtype (WT), Wilcoxon Rank-Sum test) and lowest NLR in the CH-positive patients was found in the patients with TET2 frameshift and nonsense mutations (2.95, Fig. S8).

We also sought to explore whether CH-positive patients responded differently to therapy. As the strongest neutrophil effect was seen in ER+ breast cancer, and in our dataset the majority of ER+ BRCA samples belong to patients enrolled in the MONALEESA-2/3/7 phase III trials, we investigated these patients in particular.(*22-24*) In these trials, pre- and post-menopausal women treated with Ribociclib (Kisqali) in combination with endocrine therapy (ET) showed longer progression free survival (PFS) than those treated with placebo plus ET. In these 1,503 patients, DNMT3A and TET2 CH mutations were detected at baseline in 14% and 6% of the cases, respectively (Fig. 5A). As expected, the age distribution was higher among patients with DNMT3A (p=3.6e-06) and TET2 alterations (p=0.0008) consistent with CHIP. We correlated TET2 and DNMT3A alterations with PFS to test for a predictive or prognostic relationship between CH status and response to Ribociclib. Interestingly, we found that patients with TET2 frameshift (FS) or nonsense alterations derived less benefit from Ribociclib than patients without these alterations (Fig. 5B, hazard ratio (HR) in TET2 FS/Nonsense patients: 0.96 (0.68-1.37, p-value 0.84) vs HR in TET2 WT patients: 0.58 (0.51-0.66, p-value 1.72e-15)), even after accounting for potential dose reductions due to neutropenia, the most common adverse effect of CDK4 inhibitor treatment. This differential response was not observed for patients with TET2 missense mutations or mutations in DNMT3A. These results combined with the results above point to different effects produced by different CH mutations, and provide multiple hypotheses for how a positive CH status could affect the progression and response of an oncology patient’s disease.

**Fig. 5.**
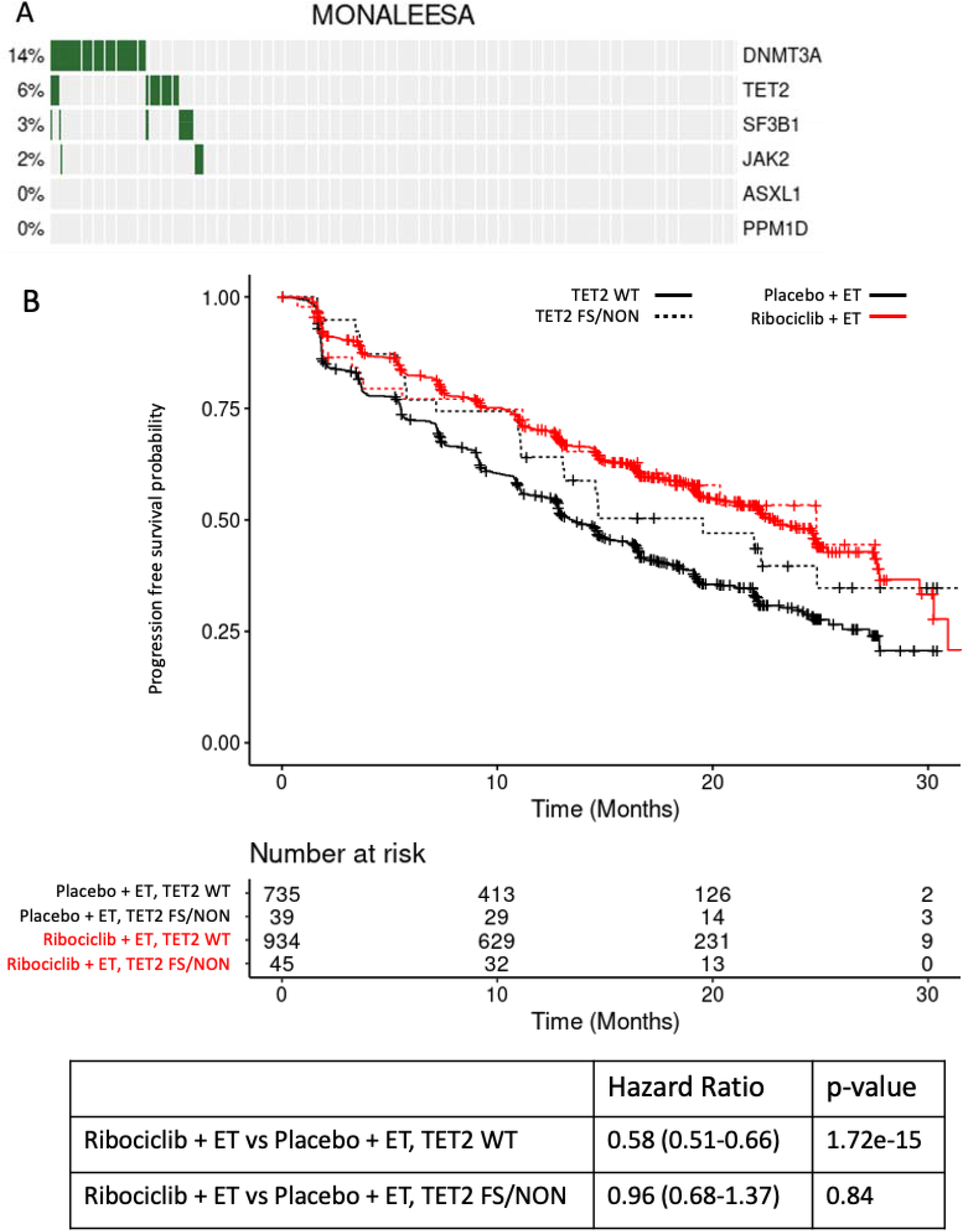
Breast cancer patients with TET2 frameshift or nonsense mutations derive less benefit from Ribociclib+ET treatment. (**A**) Incidence of canonical CH mutations in MONALEESA breast cancer patients. (**B**) Progression free survival of patients treated with ribociclib + ET (red) vs placebo + ET (black), with and without TET2 frameshift (FS) or nonsense (NON) mutations.

## DISCUSSION

Noninvasive molecular profiling of cancer patients by sequencing cfDNA from plasma samples is becoming a mainstay of modern precision medicine-based approaches in oncology. The ability to also detect CH status from these plasma sequencing assays without the need for sequencing of the matched blood sample will broaden the impact of these tests, reduce their cost by removing the need for paired WBC sequencing and give clinicians a more complete view of their patients’ health status. Here we present a machine learning method that allows for accurate detection of CH variants from plasma cfDNA without the requirement for additional sequencing of a matched blood sample.

This machine learning approach accurately distinguishes between blood and tumor derived variants using only sequence-based information and annotations derived from public databases, removing any dependency on the matched blood NGS data. Using our ML classifier on an internally generated set of cfDNA sequencing from over 4,000 patients with advanced cancer enrolled in Novartis clinical trials, we found that about 29% of patients have blood-derived mutations consistent with CH, and the incidence of these mutations increased with patient age. We also determined that the incidence of CH was similar across tumor types except for NSCLC where it was slightly higher, potentially due to higher smoking incidence. These results are consistent with earlier studies that have investigated CH in patients with solid cancers using tumor tissue and matched blood sequencing.(*2, 5*)

Observational studies on self-reported healthy individuals have found increased levels of circulating blood biomarkers of inflammation in people with CH.(*1, 16*) Here, we show for the first time, signals of inflammation both systemically and in the tumor microenvironment of CH-positive cancer patients. In CH-positive patients relative to those with no evidence of CH, we found increased expression of innate immune and inflammation pathways in the TME, as well as an increase in absolute neutrophil counts, indicating increased inflammation both locally and systemically. These data have established a relationship between CH mutation presence and inflammation, but the directionality of the relationship is not clear. Additional sequential cfDNA collections in patients may help determine the direction of this relationship. The clinical and biological impact of individual CH mutations and CH genes also remains to be determined. Here we show that nonsense and frameshift TET2 mutations are associated with poorer response to Ribociclib, but DNMT3A mutations are more strongly linked to increased NLR. These results warrant further investigation of the impact of CH status on patients’ response to cancer therapies particularly in the domain of immune oncology.

We have presented an accurate machine learning approach able to discriminate between blood-derived and tumor-derived mutations in plasma cfDNA from oncology patients. Our open-source method provides a computational solution to labeling CH variants from cfDNA, alleviating the cost-prohibitive requirement of matched WBC sequencing for cfDNA sequencing in the clinical setting. Furthermore, we used this method to characterize CH in the oncology clinical setting and correlate patient CH status with peripheral and tumor microenvironment (TME) biomarkers. Our analyses suggest that CH-positive patients show an increase in neutrophil and inflammatory activity in the TME that is reflected in changes in peripheral blood counts associated with systemic inflammation. These findings suggest that clonal hematopoiesis status might be an additional biomarker of tumor microenvironment inflammation and its impact on cancer patients’ response to therapy requires additional investigation.

## MATERIALS AND METHODS

### Model features and training

Logistic regression and random forest models were trained using 10-fold cross validation using the caret package on published data from *Razavi et al*. This data was subset to remove variants of unknown source (VUSOs), leaving 1,386 variants for training. Variants were classified as either blood-derived or tumor-derived. A table of features included in the model is provided in Table S2.

To calculate the COSMIC variant frequencies, variants from COSMIC v83 (GrCh37) were first grouped by their primary site (haematopoietic and lymphoid tissue, lung, breast, skin, kidney, liver, pancreas, stomach, large intestine, prostate, ovary, other). Freq.heme_lymph was calculated by dividing the number of COSMIC variants in haematopoietic and lymphoid tissue by the total number of variants found in “non-other” tissues. Freq.solid was calculated by summing all variants found in “non-other” tissues and non-haematopoietic and lymphoid tissues and dividing by the total number of variants found in “non-other” tissues.

### PanCancer panel

Two versions of the Novartis Institutes for BioMedical Research (NIBR) Next Generation Diagnostics (NGDx) lab’s PanCancer (PC) cfDNA assay and panel were used to generate the data analyzed here. Samples generated with PanCancer v1.0 were run using the cfDNA panel version 3-1 (N=3,140) and samples generated with PanCancer v2.0 were run using the cfDNA panel version 4 (N=1067). For both, cfDNA was extracted from approximately 4□ml of plasma (QIAamp Circulating Nucleic Acid kit, Qiagen) and then constructed into sequencing libraries with end repair, A-tailing, and PCR amplification (TruSeq Nano Library Preparation Kit, Illumina). Samples generated with PC v2.0 and panel v4 also had unique molecular identifier (UMI) adapters (custom adapters, IDT) ligated before the PCR amplification step. The constructed cfDNA libraries were then hybridized to RNA baits (SureSelect, Agilent) targeting 566 (panel v3-1) or 579 (panel v4)□cancer-relevant genes. The captured libraries were sequenced to achieve a mean unique coverage of at least 1000x using Illumina v.4 chemistry and paired-end 100-base pair (bp) reads (HiSeq, Illumina).

For samples with UMIs ligated, UMIs were trimmed from the reads using UMI-Toolkit v.1 (https://github.com/angadps/UMI-Toolkit). Reads were then aligned to the human reference genome (PC v1.0: build hg19, PC v2.0: build hg38) using the Burrows–Wheeler Aligner (BWA-MEM(*25*)). The alignments were then locally realigned and base quality scores recalibrated (GATK(*26, 27*)). For samples with UMIs, consensus reads were created using the UMI and alignment position to remove PCR-duplicate reads and sequencing artifacts (UMI-Toolkit). Single-nucleotide variants (SNVs) were identified with MuTect v.1.1.7 (*28*). Short insertion/deletion (indel) events were identified using Pindel v.1.0 (*29*). Structural variants were identified using PureCN v.1.8.1 (*30*). Chromosomal rearrangements were called using Socrates v.1 (*31*).

A position-specific error rate was calculated based on the sequencing of plasma from 24LJhealthy controls, and mutations were retained only if they had support significantly greater than the position-specific error rate. Additional potential artifacts were removed using the following filters: low allelic fraction (<0.005 unless known or probable oncogenic), poorly supported alignments (>50 MQ0 reads), low base quality (<20), low coverage (<100×) or in repetitive regions. Probable germline SNVs and indels were identified by their presence in the databases dbSNP□(https://www.ncbi.nlm.nih.gov/snp/), the Exome Sequencing Project (ESP6500SI-V2-SSA137.GRCh38-liftover, http://evs.gs.washington.edu/EVS/) and the Exome Aggregation Consortium (release 0.3; now part of gnomAD, https://gnomad.broadinstitute.org/) at appreciable frequency (ESP minor allele frequency□>0.001 or ExAC count >3 unless a known hotspot mutation). SNVs and indels were assigned a functional significance based on their presence in the Catalog of Somatic Mutations in Cancer (COSMIC v.83, https://cancer.sanger.ac.uk/cosmic) and functional effect, with mutations reported in COSMIC in five or more tumors considered as ‘known’ oncogenic, mutations with COSMIC count <5 but predicted to lead to premature truncation of the protein considered as ‘likely’ oncogenic, and all others considered to have ‘unknown’ oncogenic status. Copy number variations were considered as amplifications if the estimated copy number was ≥7, or as homozygous deletions if the estimated copy number was ≤0.5. PureCN uses a combination of the B□allele frequency of single-nucleotide polymorphisms in copy number variants and the allele frequency of somatic point mutations to determine the proportion of cfDNA derived from the tumor. The same approach was used to estimate tumor content (purity) in tumor DNA-seq. TMB was also calculated by PureCN(*30*), using the tumor content and allelic fraction information to remove germline variants and artifacts. TMB was then calculated as the number of somatic mutations per megabase of ‘callable’ coding sequence (that is, with sufficient coverage and quality).

The lower limit of detection (LLoD) in the two cfDNA panels is different (PanCancer 1.0 LLoD = 1%, PanCancer 2.0 LLoD = 0.5%). When comparing the proportion of variants classified as likely blood-derived from the two assays, more PC1.0 variants were labeled “blood-derived” than PC2.0 variants (27% vs 23%, respectively, p=6.942e-10, Fisher’s exact test).

Within canonical CHIP genes there was no difference in the proportion of blood-derived variants predicted, with 88% of PC1.0 variants classified as blood derived, compared with 89% of PC2.0 variants (p=0.71, Fisher’s exact test). When looking at the patient level, 26.6% of patients tested with PC1.0 were determined to be CH+ based on their SNV status, versus 26.4% of patients tested with PC2.0 (p=0.93, Fisher’s exact test) (Table S4).

### RNA extraction and sequencing

Sections of thickness 5□ Jµm (±1□ Jµm) were cut from all blocks received. A pathologist visually inspected archival formalin-fixed, paraffin-embedded (FFPE) slides and freshly cut slides from the tumor blocks to identify and notate the approximate percentage of tumor content in the region of interest and total tumor area (mm^2^). Depending on the tumor cell content, 4–12□ jslides were macrodissected and used for RNA isolation. If the region of interest contained <10% tumor content, further processing was canceled. RNA was coextracted from all samples available using the AllPrep RNA Extraction from FFPE Tissue Kit (Qiagen).

Ribosomal RNA from extracted total RNA was depleted using RNAseH. The rRNA-depleted sample was then fragmented, converted to complementary DNA and carried through the remaining steps of next-generation sequencing library construction—end repair, A-tailing, indexed adapter ligation and PCR amplification—using the TruSeq RNA v.2 Library Preparation kit (Illumina). The captured library was pooled with other libraries, each having a unique adapter index sequence, and applied to a sequencing flow cell. The flow cell underwent cluster amplification and massively parallel sequencing by synthesis using Illumina v.4 chemistry and paired-end 100-bp reads (Illumina).

Sequence data were aligned to the reference human genome (build hg19) using STAR v.2.4.0e (*32*). Mapped reads were then used to quantify transcripts with HTSeq v.0.6.1p1 (*33*) and RefSeq GRCh37 v.82 gene annotation.

### CH variant and patient classification

Classified blood and tumor-derived variants were further classified as CH-positive, CH-myeloid, or CH-negative. CH-positive variants were either SNVs classified as blood derived or indels classified as CHIP as described above and fell into one of six canonical CHIP genes (TET2, DNMT3A, ASXL1, JAK2, PPM1D, SF3B1). CH-myeloid variants were either SNVs classified as blood derived or indels classified as CHIP and fell into a putative myeloid driver gene, as described by *Bolton et al, 2020 (5)* (Table S5). All other variants were labeled CH-negative.

Patients were classified as CH-positive if they had at least one CH-positive variant detected. They were classified as CH-myeloid if they had at least one CH-myeloid variant detected, and no CH-positive variants detected. Finally, CH-negative patients were those with no CH-positive or CH-myeloid variants detected.

### Computational analysis and visualization

All analyses were performed in R 3.6.1 except some visualizations which required R 4.0.0. Differential expression analysis was performed using DESeq2. Plots were generated using ggplot2 and ggpubr. GO term clustering and visualization was performed using the simplifyEnrichment package. Oncoprints were generated using the ComplexHeatmap package.

## Supplementary Materials

Fig. S1. SNV classification frequency by random forest.

Fig. S2. Number of predicted CH variants per patient in CH-positive patients.

Fig. S3. Comparison of CH incidence in cancer patients and healthy controls.

Fig. S4. Odds of CH-positivity by tumor type.

Fig. S5. Log-odds ratio of CH gene incidence.

Fig. S6. Enriched GO terms in genes up-regulated in CH-positive patients.

Fig. S7. Circulating blood cell count distributions by predicted CH-status.

Fig. S8. Neutrophil to lymphocyte ratio (NLR) vs CH mutation type

Table S1. Description of training data (*Razavi et al, 2019*)

Table S2. Description of model features

Table S3. Model misclassifications on held out variants from healthy controls

Table S4. CH detection between PanCancer versions 3-1 and 4

Table S5. Putative myeloid disease driver genes (*Bolton et al, 2020*)

Table S6. Neutrophil to lymphocyte ratio by tumor type.

Data File S1. Full model training data.

Data File S2. Patient information.

Data File S3. Canonical CH patient SNVs.

Data File S4. Canonical CH patient indels.

## Acknowledgments

The authors would like to thank the patients who participated in these trials, their families and caregivers, and the staff who assisted with the trial at each site. We would also like to thank the NIBR Next Generation Diagnostics (NGDx) lab for the generation of the sequencing data. We also thank Pushpa Jayamaran and Janghee Woo for conversations relating CHIP to inflammation and Jeff Engleman, Wendy Winkler, and Audrey Kauffman for their leadership and support.

## Author contributions

Conceptualization: OAB, CDC, LF

Methodology: LF, OAB

Resources: RL, FS

Software: LF, JW, KD, CBG, NS

Investigation: LF, JW, OAB

Visualization: LF

Supervision: OAB, CDC

Writing – original draft: LF, OAB

Writing – review & editing: LF, JW, OAB, JWu, NS, CDC, OAB

## Data and materials availability

The minimal clinical and mutational data necessary to replicate the findings in the article, except those shown in Figures 4, 5, S1, and S6-8, and Table S4 and S6. Data for the excepted figures (individual trial and drug names, and non-CHIP related variants) cannot be made public to preserve patient anonymity. Raw sequencing data cannot be publicly deposited for legal and privacy reasons, as sequencing was performed for clinical purposes. Patient age is reported as a range to protect patient privacy. Code used to generate the model and figures (where underlying data is authorized for public release) is available at: https://github.com/Novartis/solid-tumor-CHIP

## Study Oversight

These studies were conducted in accordance with the principles of the Declaration of Helsinki and the Good Clinical Practice guidelines of the International Council for Harmonisation. The study protocols and all amendments were reviewed by the independent ethics committee or institutional review board at each center. All the patients provided written informed consent.

