## Supplemental materials for "Clonal hematopoiesis detection in cancer patients using cell free DNA sequencing"

Lauren Fairchild *et al.*

Corresponding author: Lauren Fairchild

Supervising authors: Catarina D. Campbell, O. Alejandro Balbin

**This PDF includes:**

Figs. S1 to S8

Tables S1 to S6

**Other Supplementary Material for this manuscript includes the following:**

Data file S1

Data file S2

Data file S3

Data file S4

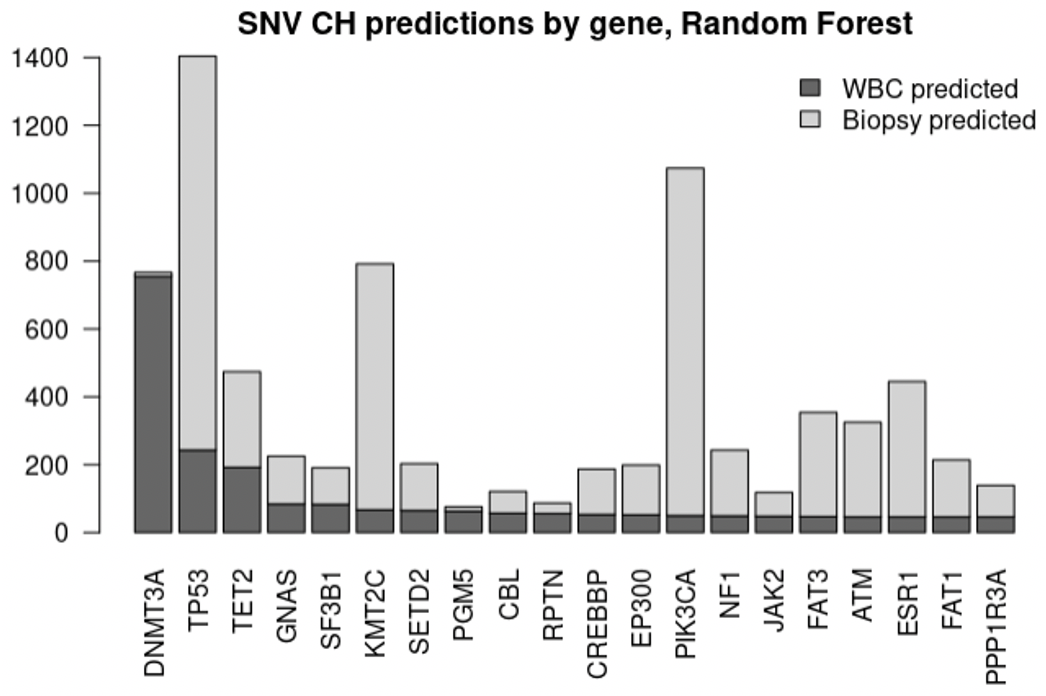

**Fig. S1. SNV classification frequency by random forest.** Proportion of SNVs within each gene classified by random forest as WBC-derived (dark grey) vs biopsy-derived (light grey).

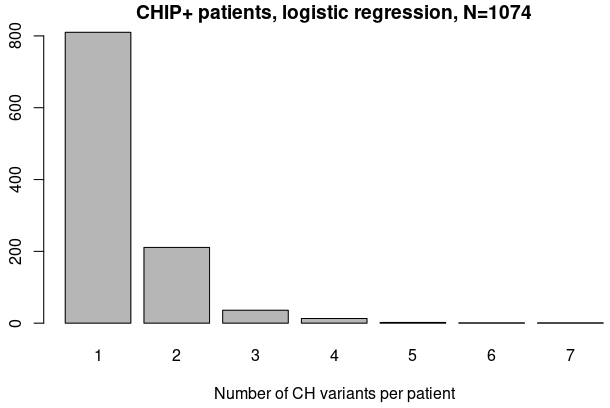

**Fig. S2. Number of predicted CH variants per patient in CH-positive patients.**

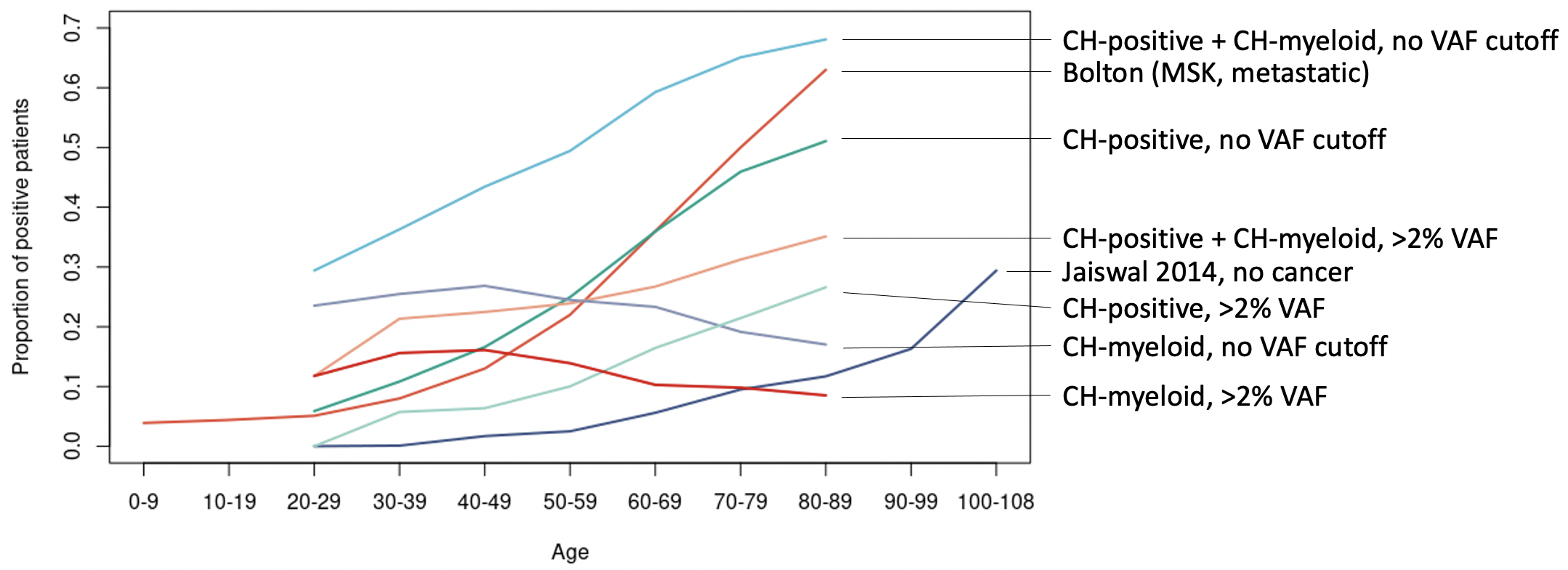

**Fig S3. Comparison of CH incidence in cancer patients and healthy controls.**

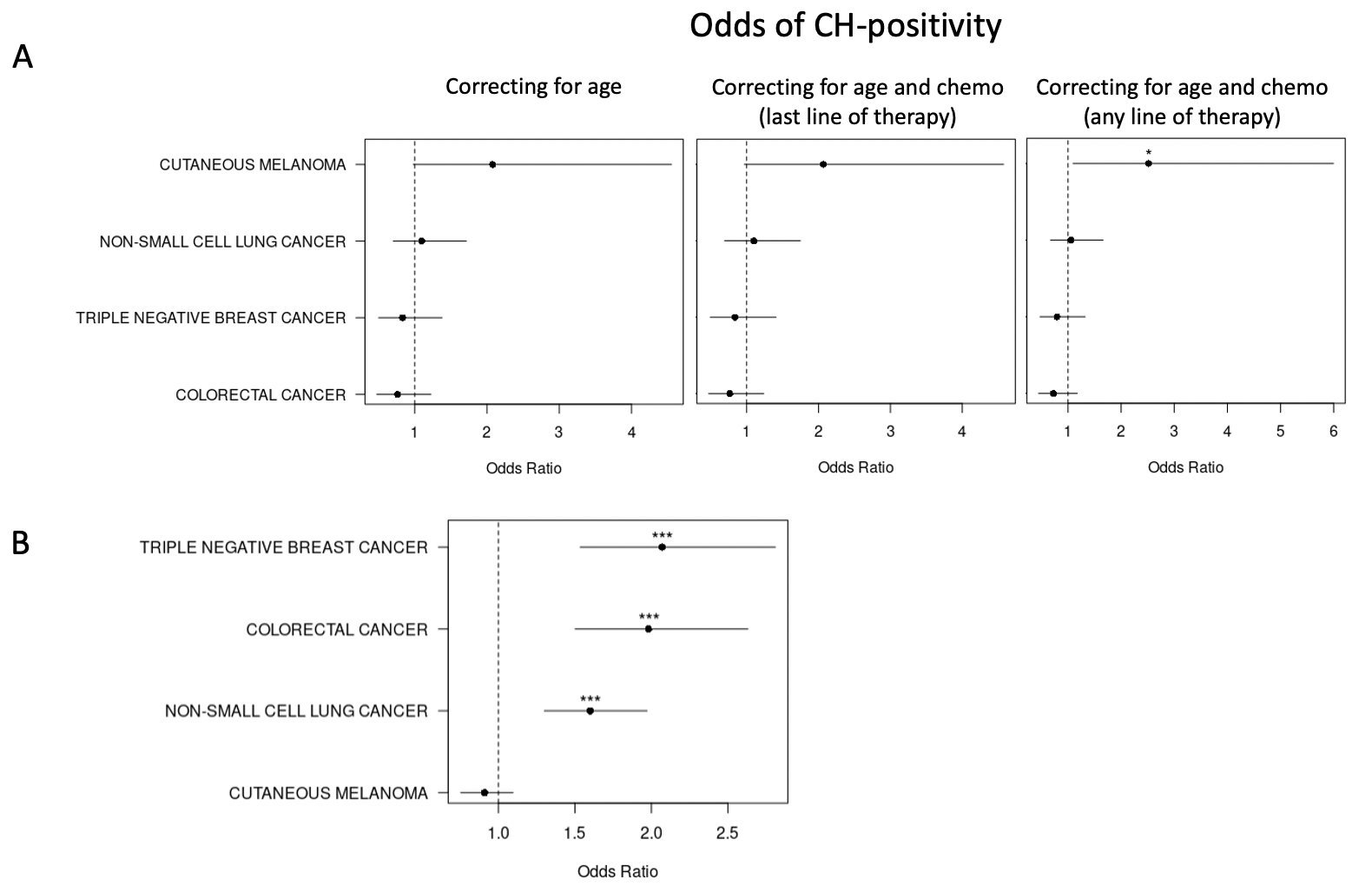

**Fig. S4. Odds of CH-positivity by tumor type. (A)** Odds of being CH-positive by tumor type relative to breast cancer for patients with prior therapy information (N=692, * indicates p-value < 0.05) **(B)** Odds of being CH-positive or CH-myeloid by tumor type relative to breast cancer, correcting for patient age. (CRC p-value: 1.79e-8, TNBC p-value: 7.43e-7, NSCLC p-value: 3.75e-10)

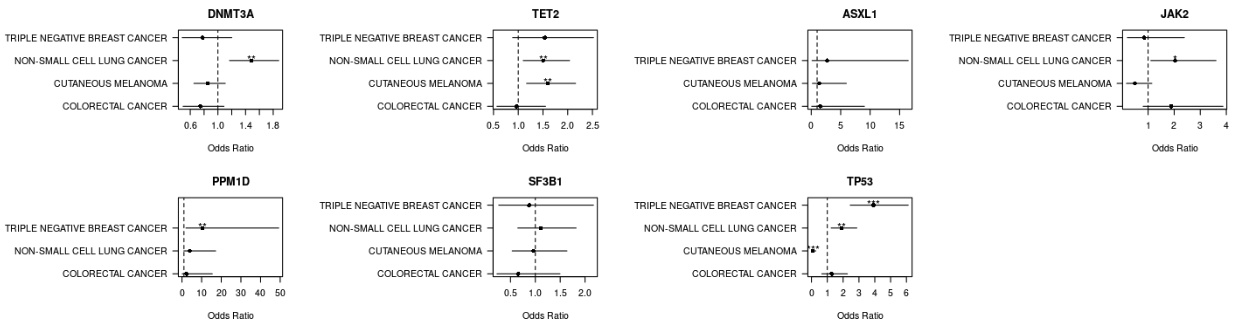

**Fig. S5. Log-odds ratio of CH gene incidence.** For each figure, ratios were calculated relative to breast cancer, correcting for patient age. (Significance codes: *** < 0.001, ** < 0.01, * < 0.05)

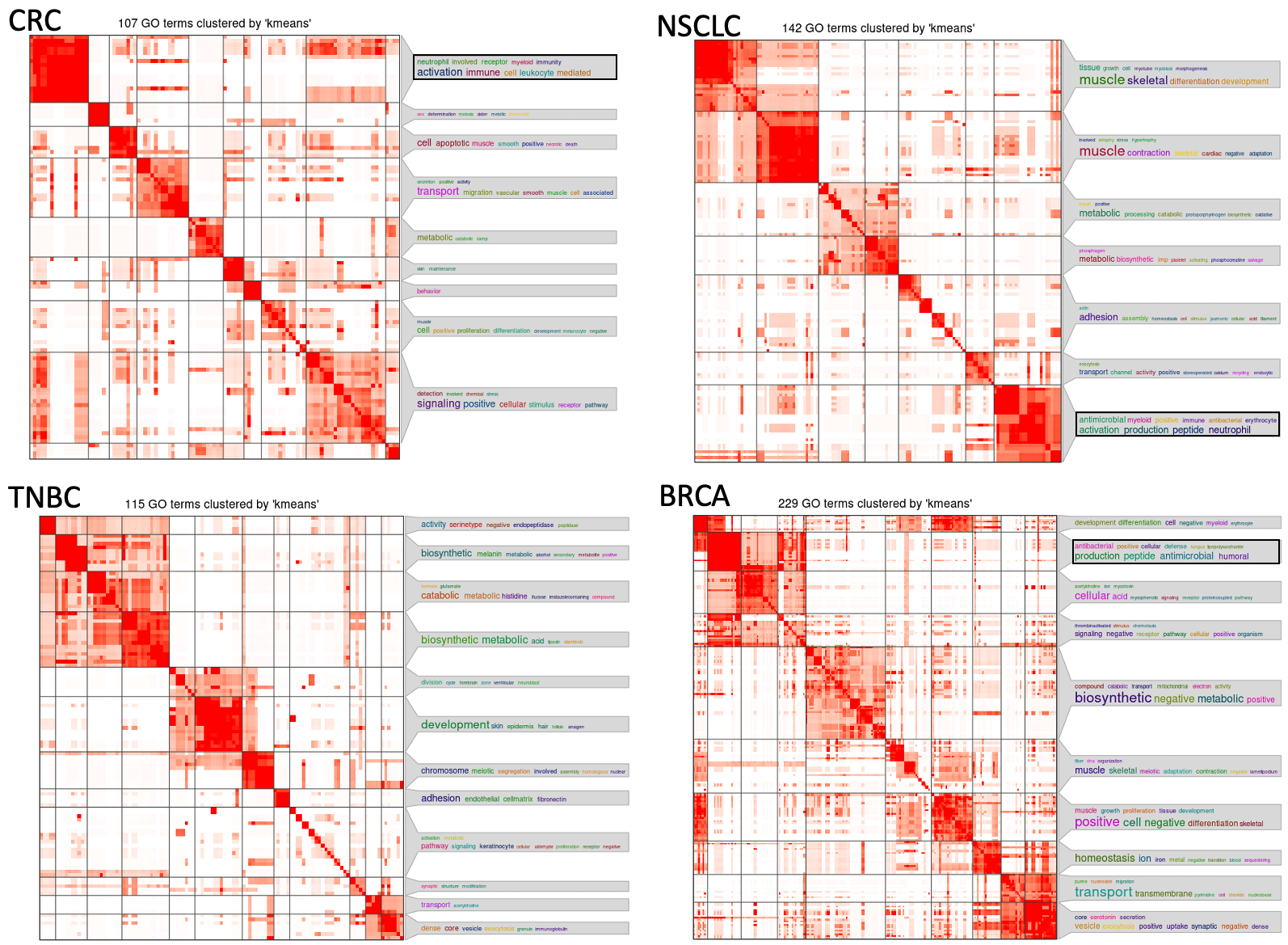

**Fig. S6. Enriched GO terms in genes up-regulated in CH-positive patients.** Each heatmap shows GO terms enriched in up-regulated genes when comparing CH-positive patients to CH-negative patients within each tumor type. GO terms are clustered using k-means clustering according to their gene membership similarity.

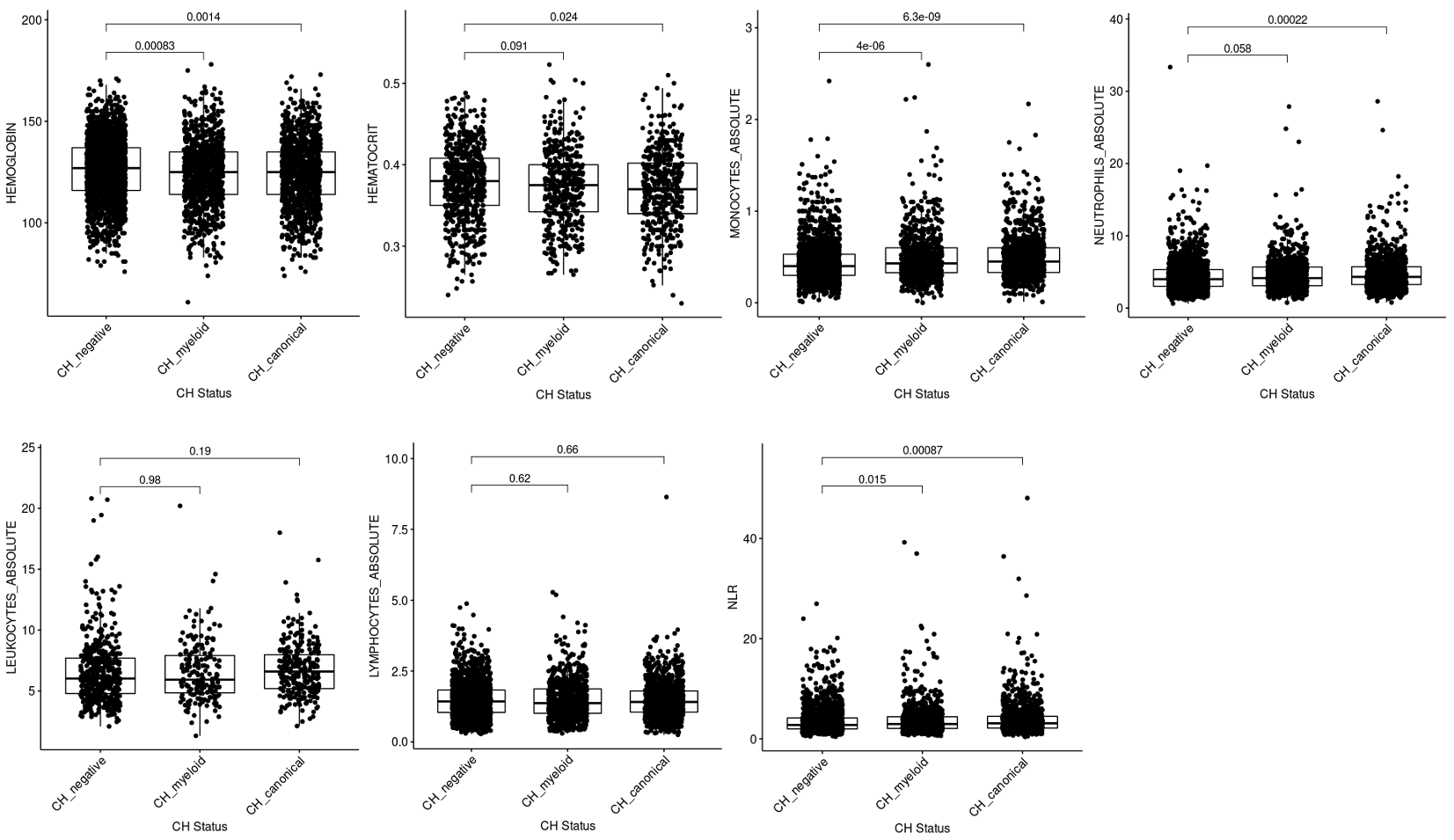

| **Medians** | **CH-negative** | **CH-myeloid** | **CH-positive** |
| --- | --- | --- | --- |
| Hemoglobin (g/L) | 127 | 125 | 125 |
| Hematocrit (L/L) | 0.380 | 0.375 | 0.370 |
| Monocytes (abs, 10e9/L) | 0.40 | 0.43 | 0.45 |
| Neutrophils (abs, 10e9/L) | 4 | 4.15 | 4.32 |
| Leukocytes (abs, 10e9/L) | 6.030 | 5.935 | 6.605 |
| Lymphocytes (abs, 10e9/L) | 1.43 | 1.37 | 1.41 |
| NLR | 2.77 | 2.97 | 3.11 |

**Fig. S7. Circulating blood cell count distributions by predicted CH-status.** NLR=neutrophil to lymphocyte ratio, t-test.

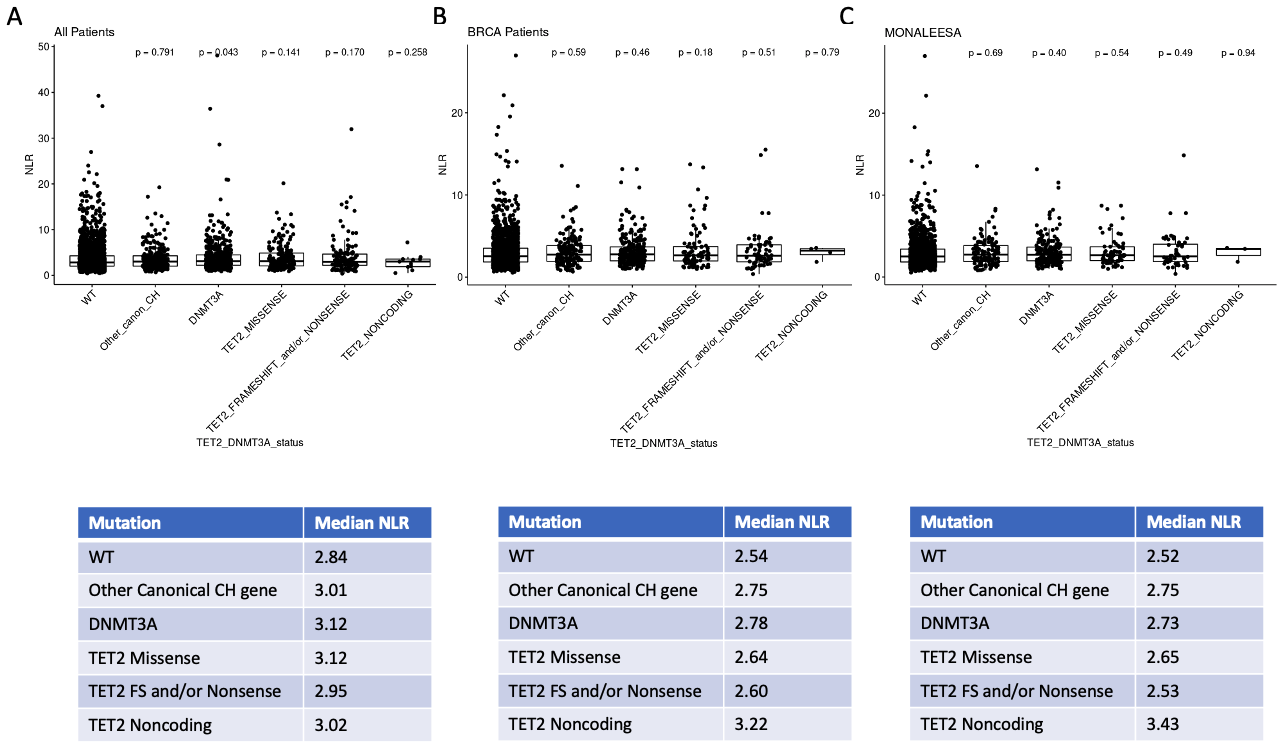

**Fig. S8. Neutrophil to lymphocyte ratio (NLR) vs CH mutation type.** For each boxplot, patients are split into those with a DNMT3A mutation, TET2 missense mutation, TET2 frameshift or nonsense mutation, TET2 noncoding mutation, other mutation in a canonical CH gene, or wild type (WT). (**A**) All patients, (**B**) Breast cancer patients, (**C**) Breast cancer patients enrolled in the MONALEESA trials.

**Table S1.** **Description of training data (*Razavi et al, 2019*).** VUSO = variant of unknown source. Note that the patients are those with at least one SNV detected.

|  | **Breast** | **Lung** | **Prostate** | **Control** | **Total** |
| --- | --- | --- | --- | --- | --- |
| N patients | 34 | 39 | 41 | 38 | 152 |
| Biopsy matched SNVs | 101 | 186 | 79 | 0 | 366 |
| Biopsy subthres SNVs | 9 | 34 | 27 | 0 | 70 |
| WBC matched SNVs | 174 | 286 | 261 | 243 | 964 |
| VUSO | 78 | 68 | 89 | 49 | 284 |

**Table S2: Description of model features**

| Feature name | Value |
| --- | --- |
| Gene name | If the variant occurs in a canonical CHIP gene or oncogene, the value is the gene name. Otherwise, “OTHER”. Gene names used: ATM, BRAF, CBL, DNMT3A, EGFR, GNAS, JAK2, KMT2C, KRAS, NF1, PTEN, RUNX1, SETD2, SF3B1, TET2, TP53. Other less common CHIP genes were grouped together under one label (BCOR, CEBPA, CREBBP, CSF1R, CTCF, CUX1, EED, EP300, ETV6, EZH2, FLT3, GATA1, GATA3, IDH1, IDH2, JAK1, JAK3, KDM6A, KIT, MPL, NPM1, PHF6, PTPN11, RAD21, STAG2, U2AF1, WT1)(*15*) |
| CH_Mutation | Classification based on the CHIP-associated somatic variants identified in *Jaiswal et al 2017* (*15*) |
| median_ctDNA | Median allele fraction of variants called within the patient |
| tq_ctDNA | Third quartile of allele fractions of variants called within the patient |
| SBS1-SBS60 | COSMIC single base substitution scores for the variant, calculated using R package deconstructSigs (<https://CRAN.R-project.org/package=deconstructSigs>), reference: signatures.exome.cosmic.v3.may2019 |
| cfdna_af | Allele fraction of the variant |
| gnomad_AF | GnomAD: Alternate allele frequency in samples |
| freq.heme_lymph | COSMIC variant frequency in heme |
| freq.solid | COSMIC variant frequency in solid tissues |
| population.frequency | ExAC: Alternate allele frequency in samples |

**Table S3. Model misclassifications on held out variants from healthy controls**

| **Gene** | **Logistic Regression Errors** | **Random Forest Errors** |
| --- | --- | --- |
| AR | 0 | 1 |
| BLM | 0 | 1 |
| BRCA1 | 0 | 1 |
| CARD11 | 1 | 1 |
| CSF1R | 1 | 0 |
| **DNMT3A** | **0** | **5** |
| EGFR | 1 | 0 |
| EPHA7 | 0 | 1 |
| GATA2 | 0 | 1 |
| INPP4B | 1 | 1 |
| IRS1 | 0 | 1 |
| IRS2 | 1 | 1 |
| KIT | 1 | 1 |
| KMT2C | 1 | 1 |
| KMT2D | 0 | 1 |
| MEF2B | 0 | 1 |
| MGA | 1 | 0 |
| MYD88 | 1 | 0 |
| NOTCH3 | 0 | 1 |
| PAK7 | 0 | 1 |
| PTCH1 | 1 | 2 |
| SDHA | 0 | 1 |
| SPEN | 1 | 1 |
| STK40 | 1 | 1 |
| **TET2** | **0** | **3** |
| TMPRSS2 | 0 | 1 |
| TP63 | 1 | 0 |
| **Total** | **13** | **29** |

**Table S4: CH detection between PanCancer versions 3-1 and 4.** Confusion matrices of logistic regression predictions for SNVs from both assay versions.

(A) Odds ratio: 0.84, p-value: 1.699e-10 (Fisher’s exact test)

| All SNVs | Biopsy | WBC |
| --- | --- | --- |
| PC v3-1 | 17544 | 6480 |
| PC v4 | 8259 | 2572 |

(B) Odds ratio: 1.09, p-value: 0.71 (Fisher’s exact test)

| SNVs in canonical CH genes | Biopsy | WBC |
| --- | --- | --- |
| PC v3-1 | 142 | 1073 |
| PC v4 | 42 | 346 |

(C) Odds ratio: 1.03, p-value: 0.78 (Fisher’s exact test)

| Patient-level CH calls (SNV+indel) | CH-positive | CH-negative |
| --- | --- | --- |
| PC v3-1 | 919 | 1498 |
| PC v4 | 325 | 544 |

**Table S5: Putative myeloid disease driver genes (*Bolton et al, 2020*)**

Boxed genes are genes considered canonical CH genes for this analysis.

| ABL1 | CDKN1B | EED | IDH1 | NOTCH1 | RRAS | WT1 |
| --- | --- | --- | --- | --- | --- | --- |
| ALK | CDKN2A | EGFR | IDH2 | NOTCH2 | RUNX1 | ZRSR2 |
| ARID1A | CDKN2B | EP300 | IRF4 | NPM1 | SETD2 |  |
| ARID2 | CDKN2C | ETV6 | **JAK2** | NRAS | **SF3B1** |  |
| **ASXL1** | CEBPA | EZH2 | JAK3 | PAX5 | SH2B3 |  |
| ASXL2 | CHEK2 | FAM175A | KDM5C | PIK3CA | SRSF2 |  |
| ATRX | CREBBP | FBXW7 | KDM6A | **PPM1D** | STAG2 |  |
| BAP1 | CRLF2 | FGFR2 | KIT | PTEN | STAT3 |  |
| BCL10 | CSF1R | FLT3 | KRAS | PTPN11 | STAT5A |  |
| BCL2 | CSF3R | GATA1 | MGA | RAC1 | SUZ12 |  |
| BCOR | CTCF | GATA2 | MPL | RAD21 | TERT |  |
| BRAF | CTNNB1 | GNAS | MYC | RAD50 | **TET2** |  |
| CALR | DICER1 | H3F3A | NF1 | RAD51 | TP53 |  |
| CBL | **DNMT3A** | H3F3B | NF2 | RB1 | U2AF1 |  |
| CDK4 | DNMT3B | HRAS | NFE2L2 | RHOA | WHSC1 |  |

**Table S6. Neutrophil to lymphocyte ratio by tumor type.** TNBC (triple negative breast cancer), BRCA (estrogen receptor positive breast cancer), NSCLC (non-small cell lung cancer), MEL (skin melanoma), CRC (colorectal cancer)

|  | NLR<3 & CH+ | NLR>=3 & CH+ | NLR<3 & CH- | NLR>=3 & CH- | p.val | Odds ratio | 95% CI |
| --- | --- | --- | --- | --- | --- | --- | --- |
| All patients | 425 | 477 | 1154 | 896 | **4.9e-06** | 1.44 | 1.23-1.70 |
| TNBC | 12 | 7 | 32 | 40 | 0.19 | 0.47 | 0.13-1.47 |
| BRCA | 265 | 211 | 824 | 450 | **5.9e-4** | 1.45 | 1.16-1.81 |
| MONALEESA | 203 | 155 | 676 | 351 | **2.2e-3** | 1.47 | 1.14-1.89 |
| NSCLC | 43 | 88 | 42 | 97 | 0.69 | 0.88 | 0.51-1.53 |
| MEL | 52 | 74 | 164 | 159 | 0.07 | 1.46 | 0.94-2.27 |
| CRC | 14 | 21 | 23 | 49 | 0.51 | 0.70 | 0.28-1.78 |
